## Supplemental text and figures for "Extrinsic Activin signaling cooperates with an intrinsic temporal program to increase mushroom body neuronal diversity"

### Supplementary information title and legends materials:

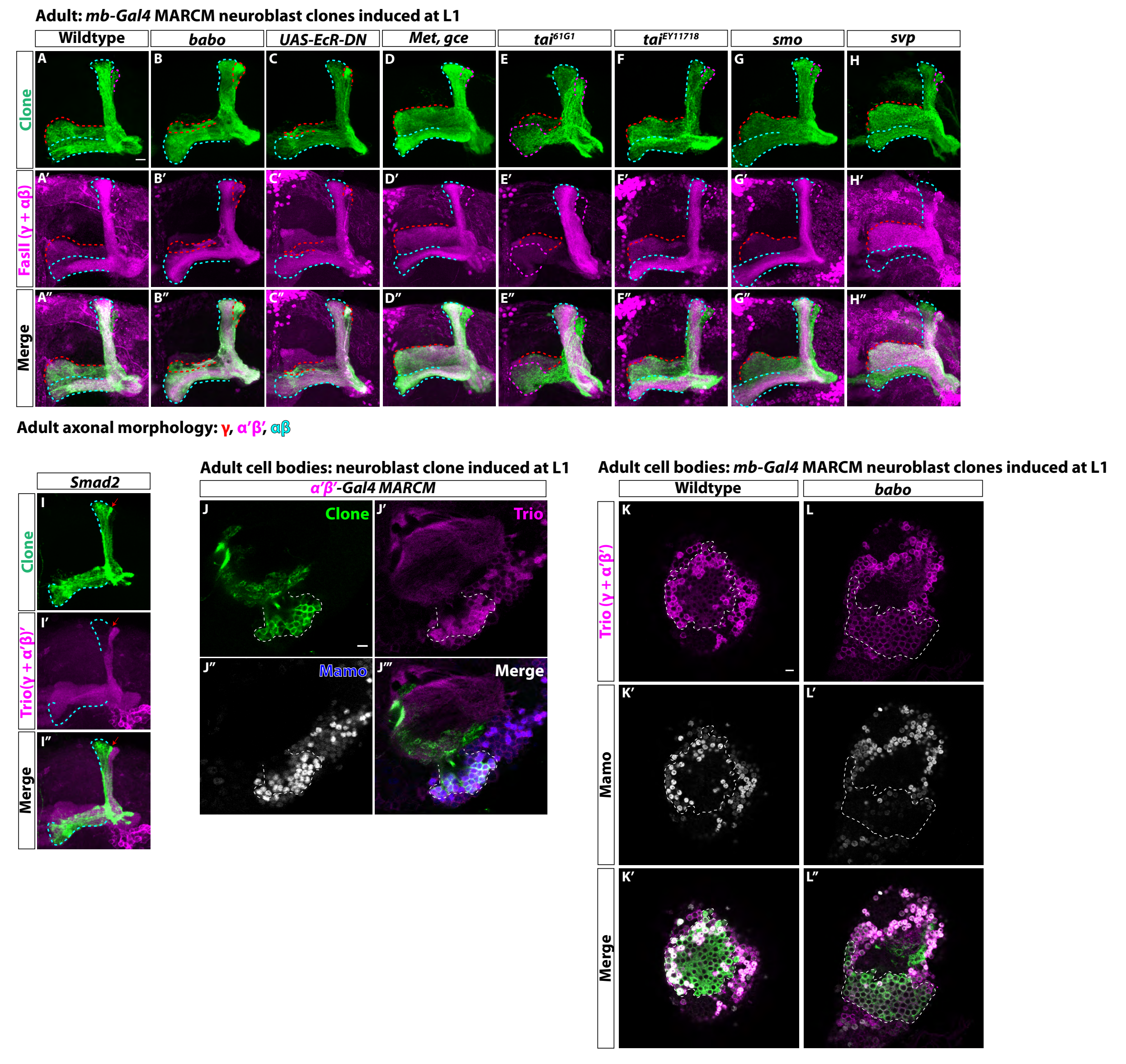


**Fig. S1. α'β' neurons are lost from the adult neuropil in Activin signaling mutant clones. A-H.** Representative images of adult mushroom body lobes from MARCM screen in which clones were induced at L1 stage. Clonally related neurons are GFP$^+$ (green). *UAS-CD8::GFP* is driven by *mb-Gal4*. All mushroom body axons, both clonal and non-clonal, are marked by FasII (γ and αβ axonal marker, magenta). Outlines mark GFP$^+$ axons, where γ axons are outlined in red, α'β' axons are outlined in magenta, and αβ axons are outlined in cyan. **A.** In wildtype clones, all three mushroom body neuronal types are present. α'β' axons are GFP$^{+}$ and FasII$^{-}$. **B.** In *babo* mutants, γ neurons do not remodel (red outline in vertical lobe). α'β' neurons are also missing, as there are no GFP$^{+}$, FasII$^{-}$ axons. **C.** *UAS-EcR-DN* expressing clones are missing α'β' axons and γ neurons do not remodel. **D-G.** No loss of neuronal diversity is observed. **H.** *svp* mutant clones (GFP$^{+}$, green) contain all three mushroom body neuronal types. **I.** *Smad2* mutant clones (GFP, green) have unpruned γ axons (red arrow) and no α'β' axons (Trio$^{+}$, magenta). **J-J'''.** Representative adult wildtype clone labelled with *R41C07-Gal4* (*α'β'-Gal4*). A single z-slice showing that strong Trio and strong Mamo label α'β' neuronal cell bodies. **K-L.** Representative, single z-slice showing strong Trio$^{+}$ and Mamo$^{+}$ cells inside and outside a GFP$^{+}$ clone in wildtype (**K**) but not in *babo* (**L**) clones. Images represent the same sections as in Figures 1F-G. Scale bars: A, 10μm; J,K, 5μm.

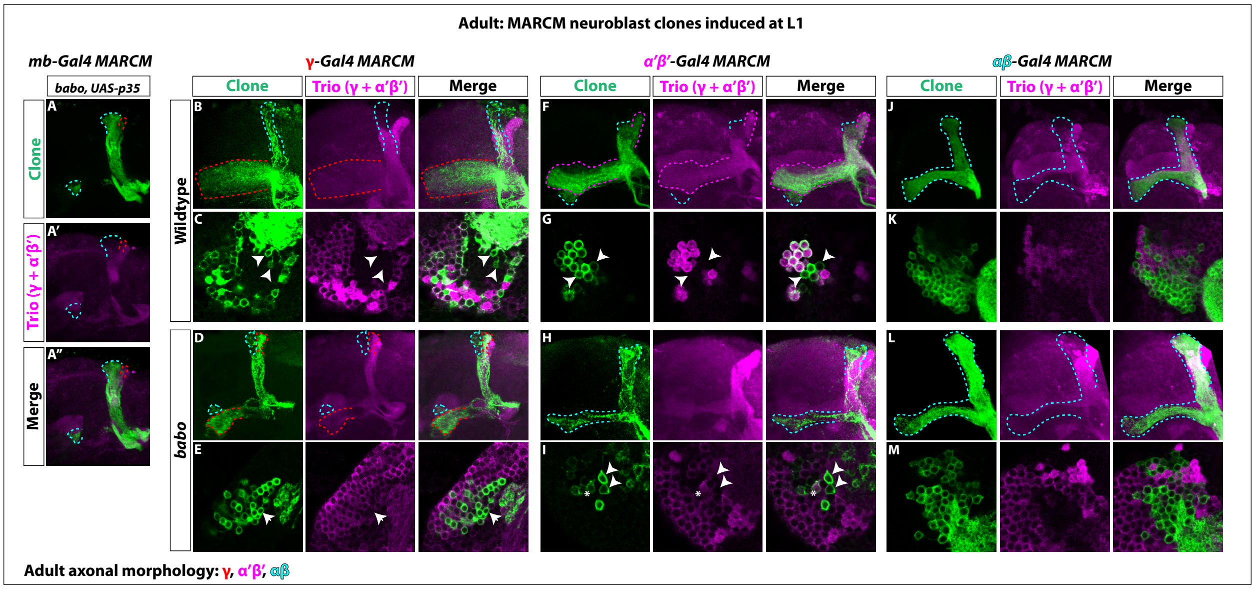


**Fig. S2. γ neuron numbers likely increase, while αβ numbers decrease, in *babo* mutant clones. A-A''.** The loss of α'β' neurons in *babo* mutant clones marked by *mb-Gal4* (GFP, green) is not rescued by blocking cell death. γ neurons (red outline) also retain their larval branching. **B-E.** Representative wildtype and *babo* mutant clones made using *R71G10-Gal4* (*γ-Gal4*) driving *UAS-CD8::GFP*. Note the presence of some $GFP^+$ axons in the αβ lobes (cyan outline). Arrowheads in the cell body region point to αβ neurons based on the absence of Trio expression. Only cells that were both $GFP^+$ and $Trio^+$ were used for quantification. **F-I.** Representative wildtype and *babo* mutant clones made using *R41C07-Gal4* (*α'β'-Gal4*) driving *UAS-CD8::GFP*. Note the presence of some $GFP^+$ axons in the αβ lobes (cyan outline). Arrowheads in the cell body region point to αβ neurons based on the absence of Trio expression. In *babo* mutants, the majority of neurons remaining are αβ ($GFP^+$ and $Trio^-$). The "*" indicates the presence of an α'β' neuron cell body based on strong Trio expression. Only cells that were both $GFP^+$ and $Trio^+$ were used for quantification. **J-M.** Representative wildtype and *babo* mutant clones made using *R44E04-Gal4* (*αβ-Gal4*) driving *UAS-CD8::GFP*.

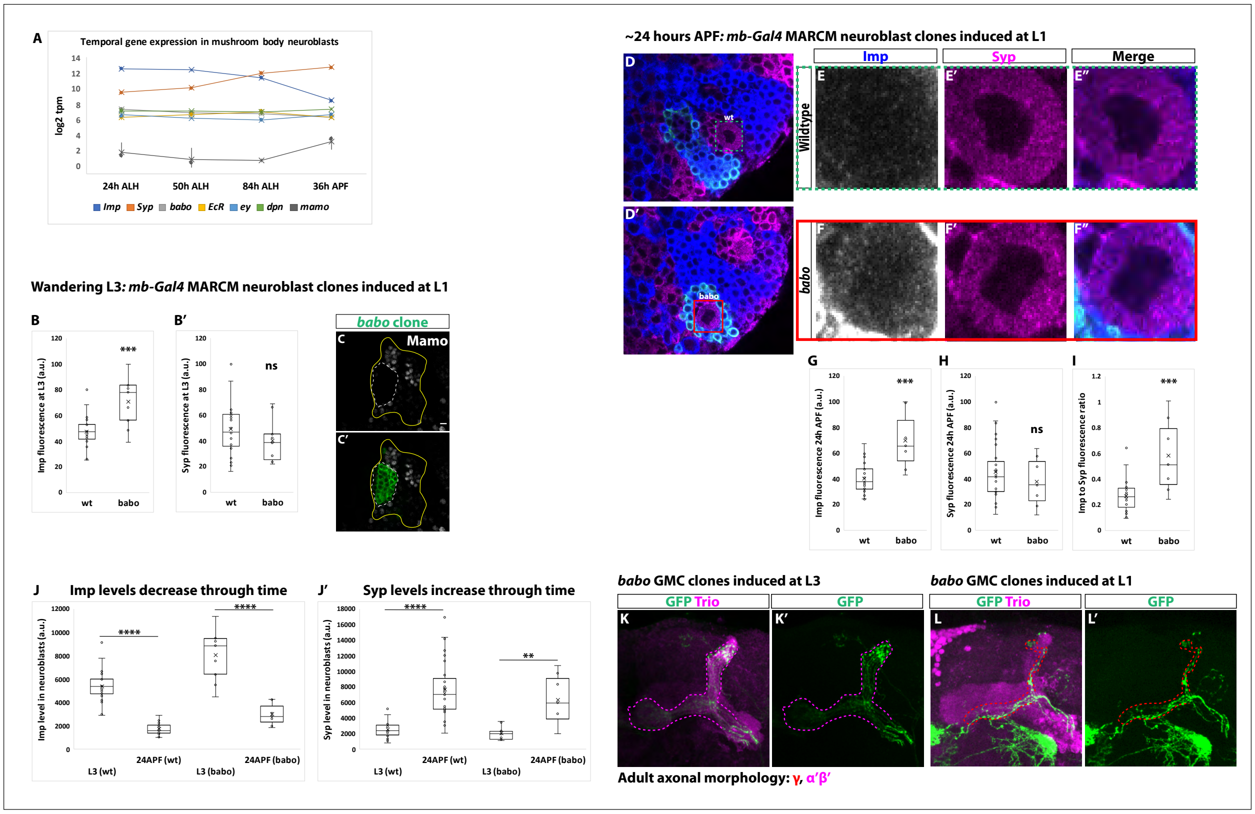


**Fig. S3. The Imp to Syp transition does not depend on Activin signaling. A.** log2 transcripts per million (tpm) of selected factors expressed in mushroom body neuroblasts through time. *babo* (gray) is not differentially expressed through time like *Imp* (blue) and *Syp* (orange). *babo* is expressed at similar levels to other important factors in mushroom body neuroblasts (i.e., *eyeless* (*ey*) and *deadpan* (*dpn*)). *mamo* served as a control since this gene is known not to be expressed in mushroom body neuroblasts. **B-B'.** Arbitrary fluorescent intensity values (scaled to 100) of Imp and Syp in wildtype (n=23) and *babo* (n=9) neuroblasts at wandering L3. The Imp level is significantly higher in *babo* neuroblasts compared to wildtype neuroblasts. Syp intensity values are not significantly different. **C-C'.** At wandering L3, *babo* mutant clones marked by *UAS-CD8::GFP* (green, white-dashed line) driven by *mb-Gal4* do not express Mamo. Mushroom body cells outside the clone (marked by Chinmo, yellow line) do express Mamo. Scale bar: 5µm. **D-D'.** Representative image of a *babo* neuroblast ~24 hours After Puparium Formation (APF) marked by *UAS-CD8::GFP* driven by *mb-Gal4* (red box) ventral to a wildtype neuroblast (green-dashed box) immunostained for Imp (blue, gray in single channel) and Syp (magenta). **E-F.** Close-up view of wildtype (**E**) and *babo* (**F**) neuroblasts from **D-D**. **G-H.** Arbitrary fluorescent intensity values (scaled to 100) of Imp (**G**) and Syp (**H**) in wildtype (n=27 neuroblasts from 6 brains) and *babo* (n=7 neuroblasts from the same 6 brains as *babo*) neuroblasts. Imp but not Syp values are significantly different. **I.** Quantification of the Imp to Syp ratio in *babo* neuroblasts compared to wildtype ~24 hours APF. *babo* neuroblasts have a significantly higher Imp to Syp ratio. **J.** Comparison of Imp fluorescent values in wildtype (wt) and *babo* mutant (babo) neuroblasts at L3 *versus* ~24 hours APF. Imp levels decrease through time independent of whether neuroblasts are wildtype or *babo* mutant. **J'.** Comparison of Syp fluorescent values in wildtype (wt) and *babo* mutant (babo) neuroblasts at L3 *versus* ~24 hours APF. Syp levels increase through time independent of whether neuroblasts are wildtype or *babo* mutant. **K-K'.** Neurons in the adult neuropil born from *babo* GMC clones induced at L3 stage and marked by *mb-Gal4* driving *UAS-CD8::GFP* (GFP, green). These neurons project axons into the Trio labeled α'β' lobes (magenta) (n=34/34). **L-L'.** In contrast, *babo* GMC clones induced at L1 show that γ neurons do not remodel in the adult neuropil, as expected (n=8/10). A two-sample, two-tailed t-test was performed. ****p<0.0001, ***p<0.001, **p<0.01, ns: not significant.

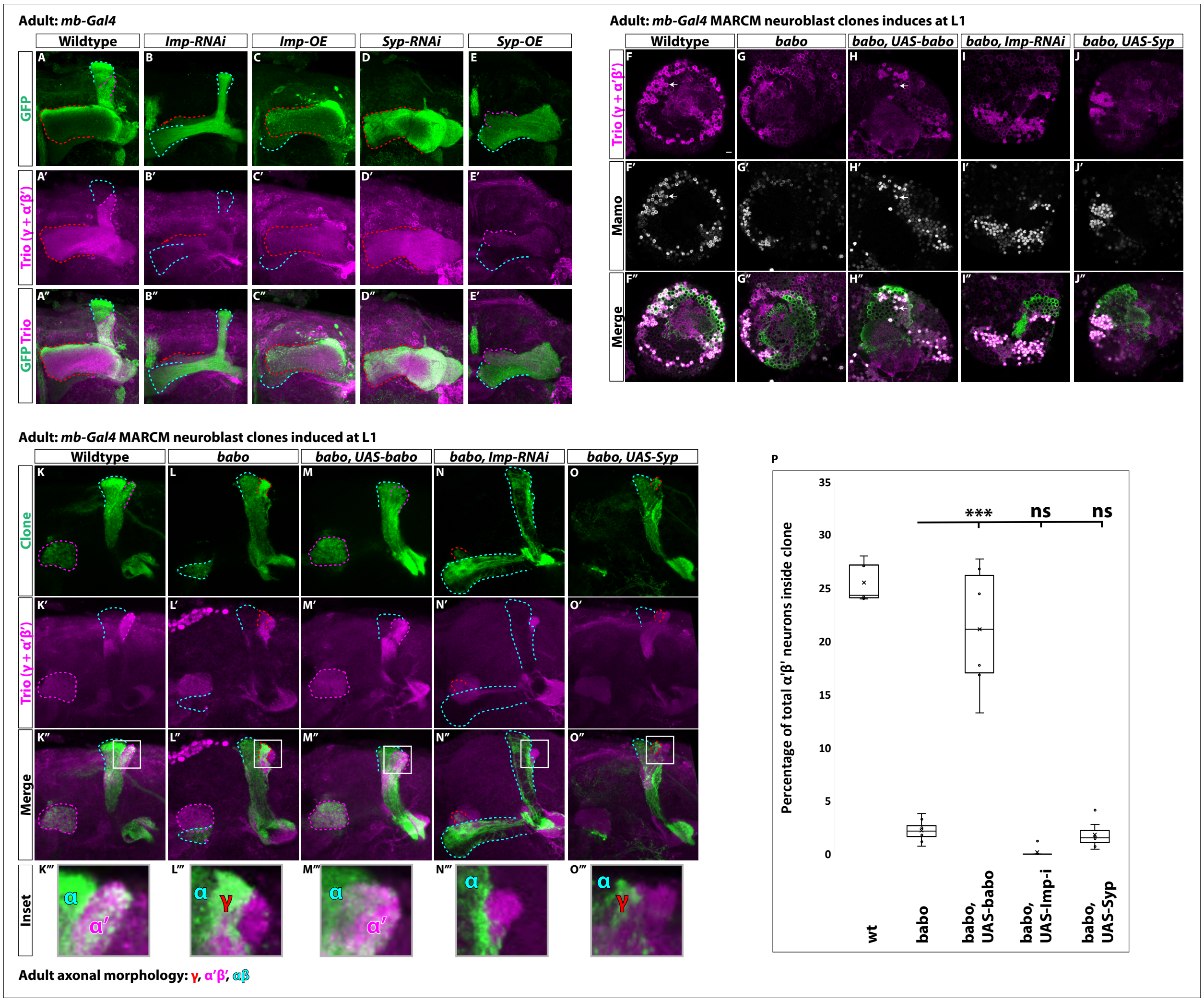


**Fig. S4. Low Imp levels are required for α'β'specification. A-E.** Representative images of adult mushroom body lobes labeled by *mb-Gal4* driving *UAS-CD8::GFP* (green) in wildtype (**A**), *UAS-Imp-RNAi* (**B**), *UAS-Imp-OE* (**C**), *UAS-Syp-RNAi* (**D**) or *UAS-Syp-OE* (**E**). When visible, γ axons are outlined in red, α'β' axons are outlined in magenta, and αβ axons are outlined in cyan. Trio (magenta) labels γ and α'β' axons. **A.** In wildtype, all three axonal types are present. **B.** With *Imp-RNAi*, the majority of γ axons (red outline) are lost and α'β' axons are completely missing. **C.** *Imp-OE* leads to mushroom body axons projecting almost entirely into the γ lobe (red outline). Some αβ axons (cyan outline) are present but are missing the vertical α projection. α'β' axons are completely lost. **D.** *Syp-RNAi* is similar but more severe than *Imp-OE* as mushroom body axons project entirely into the γ lobe (red outline). **E.** *Syp-OE* does not abolish α’β’ axons (magenta outline). αβ axons (cyan outline) are also present. However, the vertical α’ and α lobes are both missing. **F-J.** Representative images of single z slices. Strong Trio (magenta) labels α’β’ neurons while weak Trio labels γ neurons. Strong Mamo (gray) also labels α’β’ neurons while weak Mamo labels γ neurons. Arrows points to α’β’ neurons. **F.** In wildtype clones driven by *mb-Gal4*, GFP$^+$ cells (green) contain strong Trio$^+$ and Mamo$^+$ cells. **G-H.** These cells are lost in *babo* clones (**G**) but rescued in clones expressing *UAS-babo* (**H**). **I-J.** Expressing *UAS-Imp-RNAi* (**I**) or *UAS-Syp* (**J**) does not rescue the loss of α’β’ neurons. **K-O.** Representative maximum-projection images of adult mushroom body lobes from *mb-Gal4* MARCM clones induced at L1, focused on the α’ and β’ lobes. Clonally related neurons are GFP$^+$ (green). All mushroom body axons, both clonal and non-clonal, are marked by Trio (magenta). Outlines mark GFP$^+$ axons, where γ axons are outlined in red, α’β’ axons are outlined in magenta, and αβ axons are outlined in cyan. A gray box outlines the Inset panel. **K.** In wildtype, GFP$^+$ axons are observed in the α’β’ lobes (magenta outline). **L.** In *babo* mutant clones, γ neurons do not remodel (red outline) and α’β’ neurons are missing. **M.** These phenotypes are rescued by expressing *UAS-babo* inside mutant clones, as GFP$^+$ axons colocalize within the Trio labeled α’β’ lobes (magenta outline). **N-O.** Neither reducing Imp with *UAS-Imp-RNAi* or decreasing the Imp to Syp ratio by expressing *UAS-Syp* rescues the loss of α’β’ neurons. **P.** Quantification of MARCM clones represented in **K-O**. Plotted is the percentage of strong Mamo$^+$ and GFP$^+$ cells (clonal cells) versus all Mamo$^+$ cells (clonal and non-clonal cells) within a single mushroom body. The number of α’β’ neurons is quantified in wildtype (n=7, replotted from data in Figure 1H ), *babo* (n=8, replotted from data in Figure 1H), *babo*, *UAS-babo* (n=6), *babo*, *UAS-imp-RNAi* (n=7), and *babo, UAS-Syp* (n=7), In wildtype, 25.5% of the total strong Mamo expressing cells (α’β’ neurons) are within a clone while they only represent 2.2% in *babo* clones. Expressing *UAS-babo* rescues to 21.1%. In contrast, expression of *UAS-Imp-RNAi* (0.17%) or *UAS-Syp* (1.8%) is not statistically different from *babo*. Significance values were determined using a Tukey test.. ***$p<0.001$, ns: not significant.

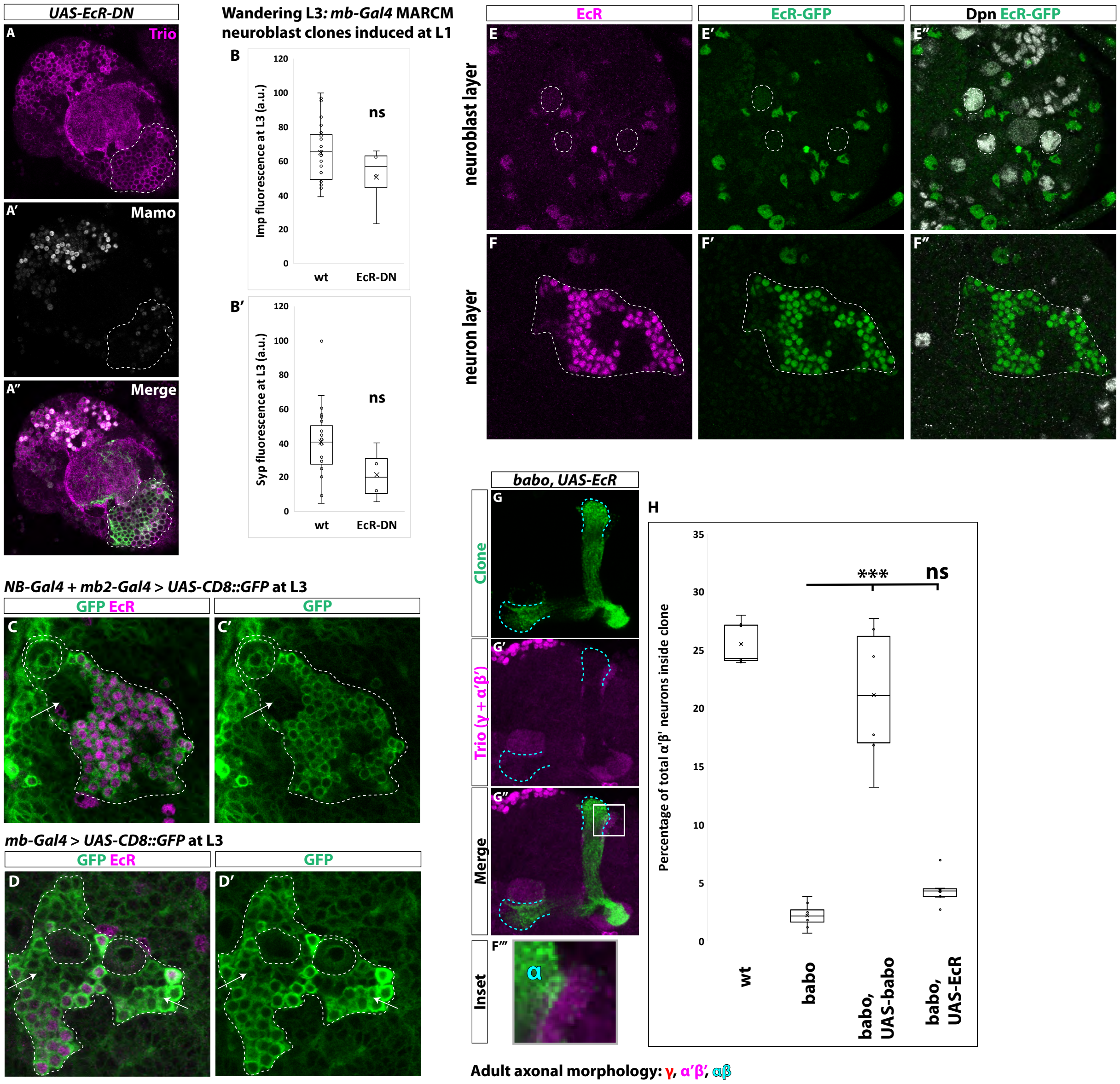


**Fig. S5. Ecdysone signaling is not necessary for α'β' specification. A.** Representative, single z-slice showing strong Trio⁺ and Mamo⁺ cells inside and outside a GFP⁺ clone expressing *UAS-EcR-DN*. Images represent the same sections as in Figure 6D. **B-B'.** Arbitrary fluorescent intensity values (scaled to 100) of Imp and Syp in wildtype (n=27) and *UAS-EcR-DN* (n=4) neuroblasts. Neither Imp nor Syp intensity values are different compared to wildtype. **C-C'.** *Insc-Gal4* + *R13F02-Gal4* (*NB-Gal4* + *mb2-Gal4*) driving expression of *UAS-CD8::GFP* at L3 stage. Strong GFP (green) is detected in mushroom body neuroblasts (white-dashed circle) but not in newborn neurons (arrows) positioned adjacent to mushroom body neuroblasts. GFP is strongly expressed in more distally positioned, mature neurons marked by EcR (magenta). **D'D'.** *OK107-Gal4* (*mb-Gal4*) driving expression of *UAS-CD8::GFP* at L3. GFP (green) is detected in mushroom body neuroblasts (white-dashed circles), newborn neurons (arrows), and mature neurons marked by EcR expression (magenta). **E-E''.** Mushroom body neuroblasts (white-dashed circles) marked by Dpn (gray) do not express EcR based on antibody staining (magenta) or an EcR-GFP (green) at the wandering L3 stage. **F-F''.** Mushroom body neurons (white-dashed outline) positioned ventrally to the mushroom body neuroblasts in **E** do express EcR but not in young neurons (black regions).**G.** α'β' neurons are not rescued in *babo* clones by expressing *UAS-EcR*. **H.** Quantification of phenotype presented in **G**. The number of α'β' neurons is quantified in wildtype (replotted from data in Figure 1H ), *babo* (replotted from data in Figure 1H), *babo*, *UAS-babo* (replotted from Figure S4P), and *babo, UAS-EcR* (n=8, 4.4%). Significance values were determined using a Tukey test.. ***$p<0.001$, ns: not significant.
